# Tactile versus Motor Imagery: Differences in Corticospinal Excitability Assessed with single-pulse TMS

**DOI:** 10.1101/2023.10.16.562530

**Authors:** Marina Morozova, Aigul Nasibullina, Lev Yakovlev, Nikolay Syrov, Alexander Kaplan, Mikhail Lebedev

## Abstract

Tactile Imagery (TI) remains a fairly understudied phenomenon despite an increased attention to this topic in recent years. Here we investigated the effects of TI on corticospinal excitability by measuring motor evoked potentials (MEPs) induced by single-pulse transcranial magnetic stimulation (TMS). The effects of TI were compared with those of tactile stimulation (TS) and kinesthetic motor imagery (kMI). Twenty-two participants performed three tasks in randomly assigned order: imagine finger tapping (kMI); experience vibratory sensations in the middle finger (TS); and mentally reproduce the sensation of vibration (TI). MEPs increased during both kMI and TI, with a stronger increase for kMI. No statistically significant change in MEP was observed during TS. The demonstrated differential effects of kMI, TI and TS on corticospinal excitability have practical implications for the development of imagery-based and TS-based brain-computer interfaces (BCIs), particularly the ones intended to improve neurorehabilitation by evoking plastic changes in sensorimotor circuitry.

**Significance Statement:** While it is known that tactile imagery (TI) engages the primary somatosensory cortex similarly to physical tactile perceptions, it is not well understood how TI affects neural processing in the primary motor cortex (M1), the area that controls voluntary movements while receiving somatosensory feedback. This study employed transcranial magnetic stimulation (TMS) to examine the responsiveness of M1 to different types of somatosensory imagery in response to TMS. TI facilitated the responses in the forearm and hand muscles but to a significantly lesser extent compared to kinesthetic motor imagery (kMI). This demonstration of the distinct effects of TI and kMI on corticospinal excitability highlights the importance of selecting an imagery strategy when using imagery to modulate cortical representation of the body. These findings have practical implications for the development of imagery-based brain-computer interfaces (BCIs) intended for rehabilitation of sensorimotor impairments.

## Introduction

Kinesthetic and tactile imagery are subdomains of the mental imagery, which is a simple but practically significant mental endeavor where actions and perceptions are imagined but not performed overtly. Most of us can imagine motor movements like bending a finger or playing the piano. This kind of motor imagery usually has both visual and kinesthetic components, that is movements are mentally visualized and also reproduced as feelings in the body parts and muscles. The latter component, which we refer to as kinesthetic motor imagery (kMI), is associated with an activation of the same neural structures that participate in the control of overt motor activity (1–3). kMI is used in sports as a way to rehearse and improve motor performance (4). Additionally, kMI is utilized as therapy (5) and neurorehabilitation approach (6).

By contrast to kMI which has been studied for decades, a somewhat different type of somatotopic imagery which we call tactile imagery (TI) has received much less attention. During TI, subjects imagine experiencing sensations arising from the skin, such as being touched with a brush or stimulated with a vibrating probe. We assert that the distinction between kMI and TI is important because kinesthetic and tactile imagery engage different types of somatosensory modalities (proprioceptive versus cutaneous) as well as incorporating imagery of different types of actions (self-initiated movements versus passive application of stimuli). Indeed, kMI involves a mental enactment of kinesthetic sensations resulting from voluntary movements (muscle contraction, joint position, tendons tension, *etc*.) and in some versions the sense of body position (2,7–9) whereas TI is about perceiving touch in a particular location on the skin surface (10–12). Expectedly, TI specifically activates the primary somatosensory cortex (10,11,13,14), as well as the areas generating the act of mental imagery, namely the prefrontal cortex (11) and parietal cortex (15). The TI-induced cortical activities can be revealed using electroencephalographic (EEG) recordings, where event related desynchronization (ERD) of the mu rhythm is the most prominent effect (12,16,17).

While EEG-based BCIs could be considered a practical goal of the research on kMI and TI (17–19), EEG recordings alone are insufficient for further development of such BCIs. Clearly, additional approaches are needed to study cortical processing under different conditions of imagery in greater detail (20,21). A variety of recording and stimulation methods could provide valuable insights. Here we focused on the effects of different types of imagery on corticospinal excitability assessed with the use of transcranial magnetic stimulation (TMS). Corticospinal excitability, measured using TMS, is known to change during the performance of sensorimotor related tasks such as kMI and action observation (7,22–25), which makes TMS an appropriate tool to study the effects of TI, as well. As to the effects on spinal excitability, kMI and TI-induced changes in the F-wave were previously reported (26).

In the present study, we applied vibrotactile stimuli to the participants’ fingertips (i.e. tactile stimulation, TS) and then asked the participants to imagine experiencing these sensations (TI) or, in a different set of trials, to imagine performing finger taps (kMI). During the performance of these tasks, TMS was applied to the hemisphere contralateral to the examined hand, which resulted in motor-evoked potentials (MEPs) in the forearm and hand muscles. We found that MEPs differed depending on the stimulation/imagery conditions. Namely, MEPs were stronger during kMI than during TI. This finding indicates that although both types of imagery involve the somatosensory domain, they affect corticospinal excitability in different ways. The possibility of manipulating imagery to evoke differential effects on the connectivity between M1 and the spinal cord opens new directions for future research and practical applications.

## Methods

### Participants

Twenty-two healthy right-handed volunteers (4 females and 18 males, 24.5±3.3 years old; mean±standard deviation) participated in the study. The participants had 6–9 hr of sleep prior to the TMS procedure, no alcohol or medication 24□hr before the procedure, and no coffee intake in the preceding 2 hr. All subjects were screened for contraindications to TMS (27). The experimental protocol was approved by the Institutional Review Board of the Skolkovo Institute of Science and Technology and followed by the Declaration of Helsinki. All participants gave their informed consent to get involved in the study.

### Experimental Design

In the description of our experimental design (Fig 1A), we used the recommendations of the OHBM COBIDAS MEEG committee for reproducible EEG and MEG research (28). Each participant participated in one experimental session lasting approximately 1.5-2 hours. The preparatory procedures included detailed explanations, instructions, and the completion of informed consent forms.

**Figure 1.**
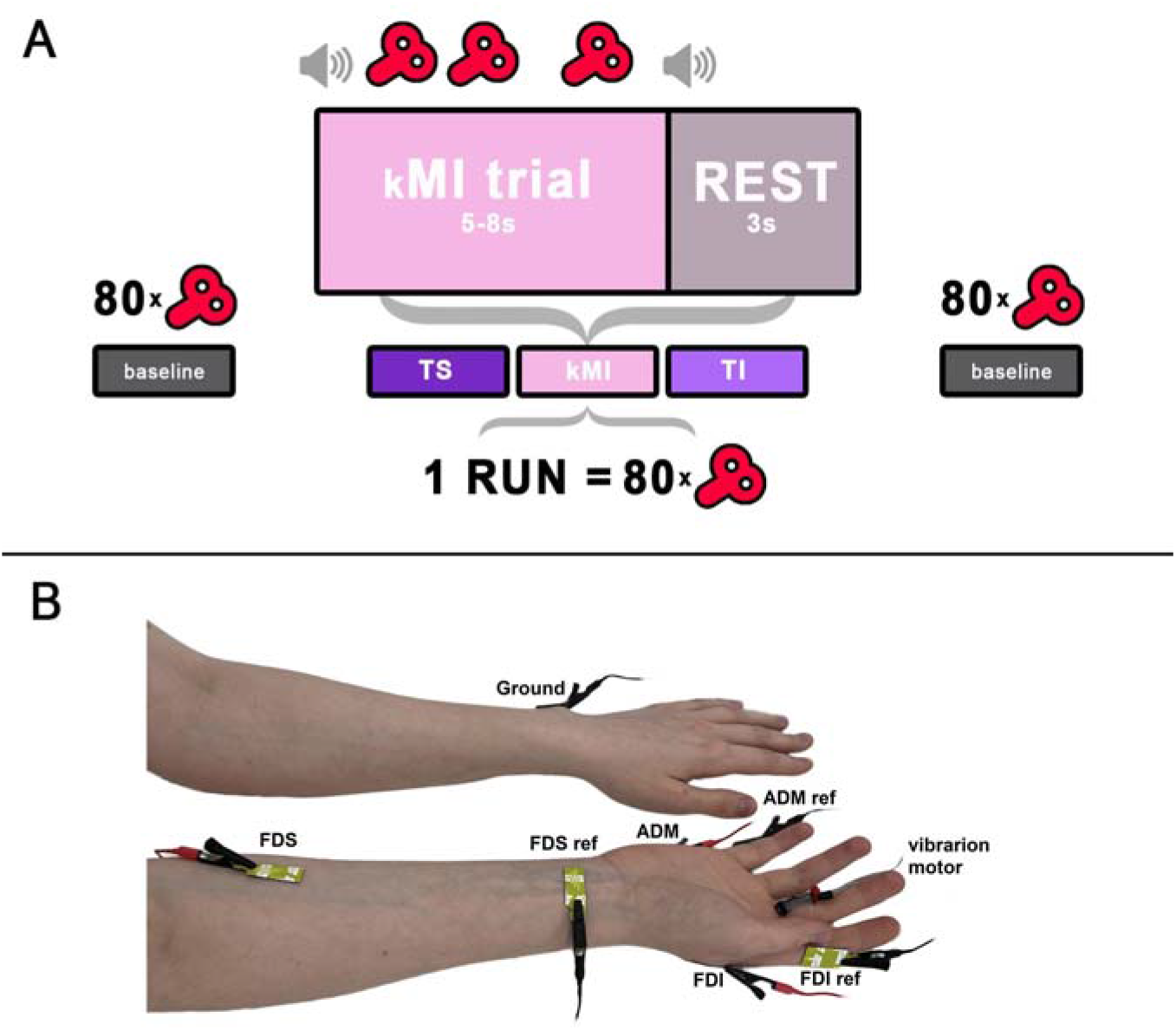
Experimental design. **A**. Schematics of task sequence. Baseline state MEP measurements were conducted before and after the session. The task included three experimental conditions (TS, kMI and TI) performed in separate runs. Each run consisted of several trials lasting 5-8 s of a period of task performance followed by a 3-s period of resting. The total number of TMS pulses was 80 for each condition. **B**. Placement of the EMG electrodes and vibratory motor on the forearm and hand.

Following the preparatory procedures, the participant was seated in a comfortable TMS-chair in front of the television screen (Samsung 46″ TV, South Korea). The screen was placed at a distance of 2.5 m from the participant. Several runs corresponding to each condition (baseline, TS, TI, and kMI) were performed. Participants were asked to fix their gaze on a cross displayed on the screen during all runs.

During the baseline runs (conducted before and after the task performance), TMS pulses were delivered to the left M1 while subjects were seated, fully relaxed, and not performing any task. The first baseline run was followed by three experimental runs ordered by random selection across subjects. Each of these runs consisted of consecutive trials during which the participants performed the required task. Trial duration was 5-8 s, and the intertrial interval was 3s.

The imagery trials began with an audio-verbal cue “imagine”. The TS trials began with the audio-verbal cue “feel”. The task performance on the trials was terminated with a beep instructing participants to stop performing the task and rest. During each trial, 2-3 randomly timed TMS pulses were applied to the left M1 with an interpulse interval of not less than 2 s. EMG activity was monitored during each trial to control for muscle activity and avoid delivering TMS during the periods of increased EMG activity.

Each run ended after a total of 80 TMS pulses were applied. Because of these settings, the number of trials per condition varied from 30 to 40. After the imagery and TS tasks were completed, the second baseline run completed the session.

Before the TI run, the participants were acquainted with the tactile sensations produced by vibration of proximal parts of the index, middle, ring and little fingers. Additionally, we evoked tactile sensations with a brush of these fingers and the distal part of the palm. During the TI run, the participants were asked to mentally reproduce the tactile sensations as emerging from these skin locations (12,16). The participants were instructed not to look at the hand and instead direct their gaze to the fixation cross. During the TS task, the participants followed the instruction to attend to their sensation of vibrations applied to the base of the middle finger. During the kMI task, the participants mentally reproduced the kinesthetic sensations associated with performing finger movements. The finger movements were flexions and extensions at the metacarpophalangeal joint – the action often called finger tapping. We chose this specific type of movement based on our previous work, where participants learned it with ease and comfortably performed it repetitively (20,21). Prior to the kMI run, the participants practiced performing finger movements at various speeds and executing isometric contractions and memorized the kinesthetic sensations associated with the movements, including muscle contraction, tendon tension, and joint position. The imagined finger movements were to be performed in a random order (21).

### EMG

Surface EMG recordings were conducted using bipolar probes placed over the right *m. flexor digitorum superficialis* (FDS), *m. first dorsal interosseous* (FDI) and *m. abductor digiti minimi* (ADM). The active electrode was placed over the muscle belly, the reference electrode over the tendon distally, and the ground electrode was placed on the wrist of the opposite (left) hand (Fig 1B). EMG data were sampled at 3 kHz using the built-in Nexstim EMG system (Nexstim Ltd., Helsinki, Finland).

### TMS

Single-pulse, biphasic TMS pulses were applied to the left primary motor cortex (M1) using a cooling co-planar figure-of-eight induction coil (d=70□mm; Nextim Ltd., Finland) connected to an eXimia magnetic stimulator (Nexstim, Finland, v3.2.2). The resting motor threshold (RMT) for the FDS muscle was determined at the beginning of experimental session using the Nexstim’s built-in threshold detection system as the minimum stimulation intensity (% of maximum stimulation output) required to elicit peak-to-peak MEP amplitudes in the FDS muscle that exceeded 50 μV in 5 out of 10 consecutive pulses (29,30). During the experimental session, the stimulation intensity was 110% of the RMT. The coil orientation was kept constant using the eXimia NBS navigation system (Nexstim, Finland). Coil tracker accuracy (3D rms error) was ±2 mm for the position and ±2° for the angle.

### Vibrotactile stimulation

TS was delivered using a custom designed vibrotactile stimulator based on Arduino UNO (Arduino, Italy) synced with stimulus presentation software. A flat vibratory motor (d=6 mm; max speed of 12.000 rpm) was placed on the middle finger of the right hand. Variable-frequency stimulation patterns were used: stimulation was delivered as 100-ms pulses with vibratory frequency selected in the range 3.000-12.000 rpm, with the inter-pulse intervals varying in the range 200-400 ms. This random stimulation pattern was used to reduce tactile habituation and residual tactile sensations (12,16).

### Data Analysis

The data analysis started with splitting the EMG records into the epochs timed -0.5 to 0.1 s relative to the occurrence of the TMS pulse. These epochs were then baseline corrected by subtracting the mean for the -0.5 to 0 s window. Next, we calculated the peak-to-peak EMG amplitude by computing the difference between the maximum and minimum values within the 10 to 50 ms temporal window relative to the stimuli onset. We then calculated the median MEP peak-to-peak amplitude for each participant and each condition (the baselines, kMI, TI, TS).

To account for the inter-individual variability, we percent normalized the median peak-to-peak amplitude for the kMI, TI and TS conditions to the baseline MEPs collected at the beginning of the experimental session:

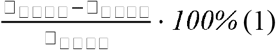

Positive values of this metric corresponded to a MEP increase from the baseline and negative corresponded to a decrease (Fig 2). The metric was calculated for each participant, and then statistical analyses were conducted for the sample of participants. For pairwise comparisons, we employed the Wilcoxon signed rank test. To assess the across-participants correlations between the MEP amplitudes in different conditions, we used Spearman’s rank correlation coefficient.

**Figure 2.**
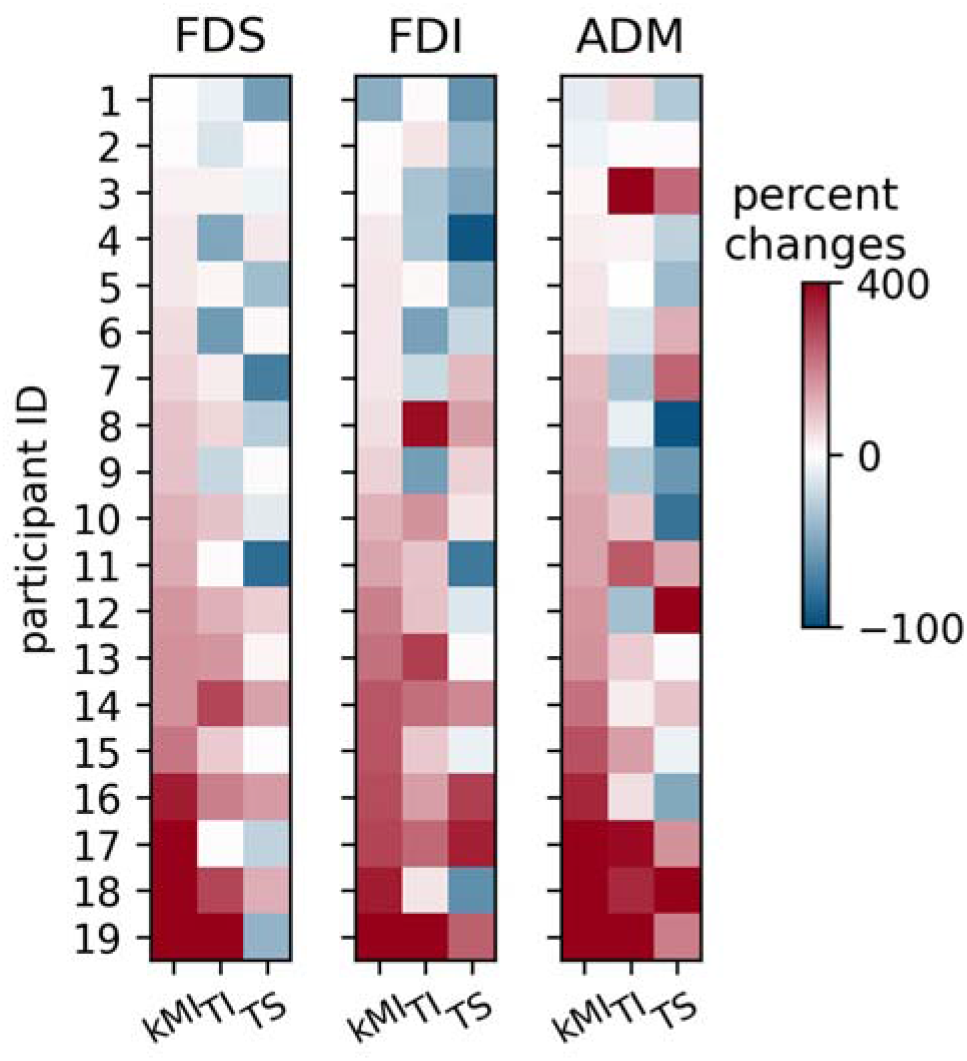
Median MEPs for each of 19 participants (color coded). The values represent changes (as percentage) from the baseline preceding the task performance. Data are shown for the kMI, TI and TS conditions and for 3 muscles: FDS, FDI and ADM.

Additionally, we used a one-sample Wilcoxon test to determine whether the normalized median peak-to-peak amplitudes were different from zero. This test determined whether there were significant changes in the MEP amplitude compared to the baseline.

Three participants were excluded from the analysis because they reported being unable to perform the required imagery. Notably, their MEP decreased when they attempted kMI. Yet, even if these participants were kept, none of the statistical results described below changed.

## Results

### General changes in MEP amplitudes

Using the Wilcoxon signed rank test for paired samples, we compared the baseline-normalized MEP amplitudes across the kMI, TI and TS condition for each muscle (Fig 3A, Fig 3B). We found a statistically significant increase in the MEP amplitude for the FDS muscle during kMI as compared to the TI and TS conditions (*W*_*kMI-TI*_ = 31, *p*_*kMI-TI*_ = 0.0082; *W*_*kMI-TS*_ = 2, *p*_*kMI-TS*_ < 0.0001). The FDS MEP was significantly stronger during TI than TS (*W* = 25, *p* = 0.0033). We found a similar tendency for the MEP to increase during kMI compared to TS in the other two recorded muscles (FDI, ADM), with some of these effects being statistically significant. Namely, the FDI response was stronger during kMI and TI than during TS (*W*_*kMI-TI*_ = 44, *p*_*kMI-TI*_ = 0.0401; *W*_*kMI-TS*_ = 29, *p*_*kMI-TS*_ = 0.0062), and the ADM muscle response was stronger during kMI than during TI (*W* = 35, *p* = 0.014).

**Figure 3.**
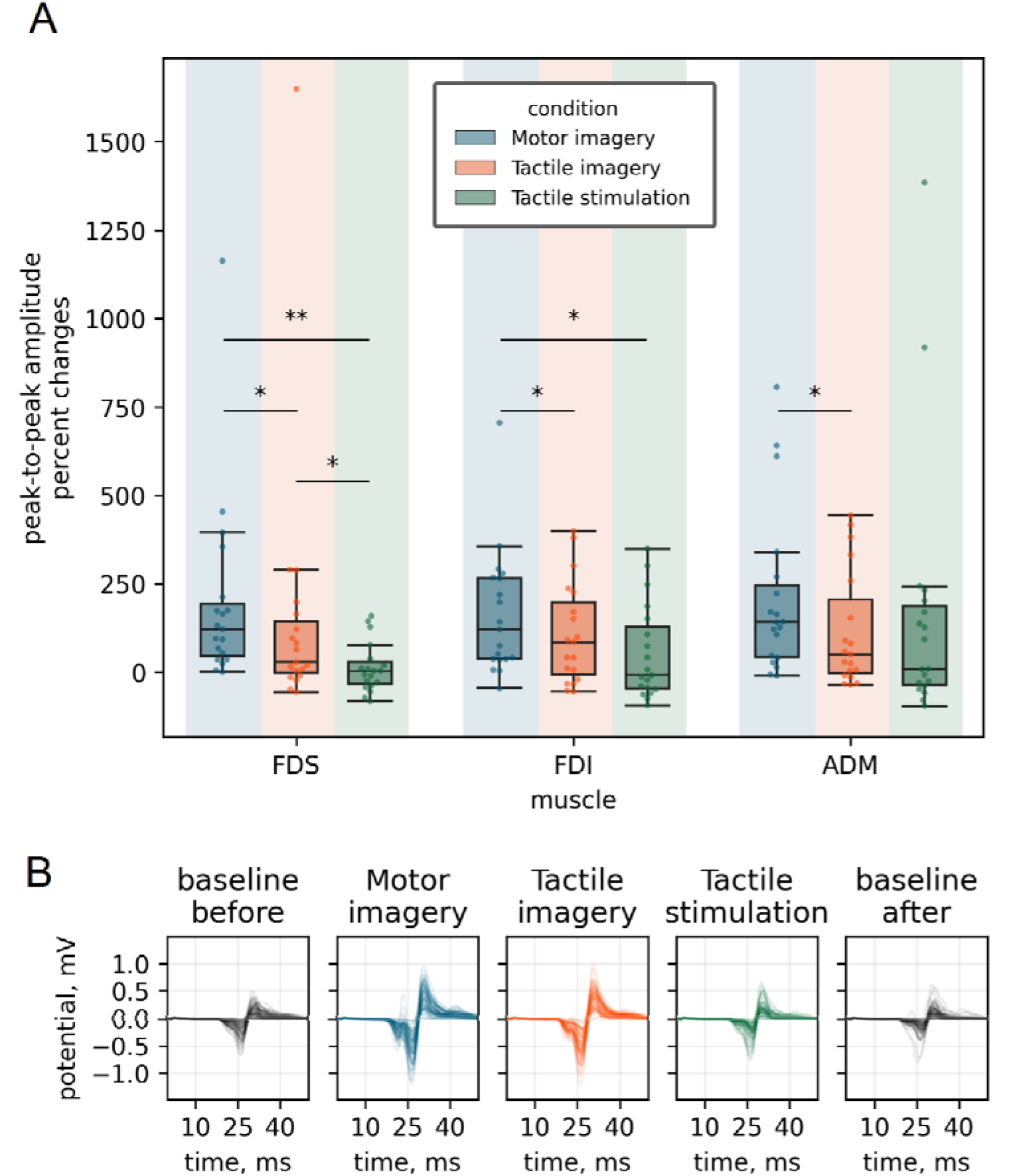
MEPs across the task conditions. **A**. Median peak-to-peak normalized amplitudes across the conditions. Data are shown for 3 muscles. Wilcoxon signed rank test was used for pairwise comparisons between conditions across the kMI, TI and TS conditions. Asterisks denote the statistically significant differences. **B**. MEP traces for the FDS in a representative participant.

Using a one-sample Wilcoxon test, we analyzed whether the changes in MEP amplitude from the initial baseline were statistically significant. This analysis was performed for the kMI, TI, TS conditions and the last baseline measured after the task performance. Increases in MEP were statistically significant for the kMI, TI and the last baseline for all muscle, namely FDS (*W*_*kMI*_ = 0, *p*_*kMI*_ < 0.0001; *W*_*TI*_ = 32, *p*_*TI*_ = 0.0095; *W*_*baseline after*_ = 30, *p*_*baseline after*_ = 0.0071), FDI (*W*_*kMI*_ = 7, *p*_*kMI*_ < 0.0001; *W*_*TI*_ = 29, *p*_*TI*_ = 0.0062; *W*_*baseline after*_ = 24, *p*_*baseline after*_ = 0.0028), and ADM (*W*_*kMI*_ = 3, *p*_*kMI*_ < 0.0001; *W*_*TI*_ = 31, *p*_*TI*_ = 0.0082; *W*_*baseline after*_ = 43, *p*_*baseline after*_ = 0.0361). Additionally, for the FDS and FDI, we found a statistically significant correlation between the MEP amplitudes during the kMI and TI conditions (Spearman’s correlation coefficient, *r*_*S*, *FDS*_ = 0.71, *p*_*FDS*_ = 0.0006; *r*_*S*, *FDI*_ = 0.65, *p*_*FDS*_ = 0.0025).

Additionally, to test for the presence of somatosensory receptor adaptation during the TS condition, we compared the median amplitudes for the first and last quarters of the TS runs using the Wilcoxon signed rank test. We did not find any statistically significant differences (*W* = 66, *p* = 0.258).

During the preparation for the TS session, we also asked the participants about the intensity and skin location of their sensations of vibration. According to the participants, these sensations were not limited to the stimulated finger but were also felt in the proximal parts of index, middle, ring and little fingers and in the distal part of the palm. These parts of the hand were located in proximity of the FDS muscle hotspot and, because of this location, were attended to during the kMI task.

### Baseline before and after the experimental session

To determine whether the sensorimotor tasks participants performed during the session affected baseline motor excitability, we compared MRP amplitudes from the first and last baseline runs. There was an increase in median MEPs amplitudes for baseline recordings after the experimental session, but these effects were statistically significant only in the FDS (*W* = 30, *p* = 0.0071) and FDI (*W* = 36, *p* = 0.016) (Fig 4).

**Figure 4.**
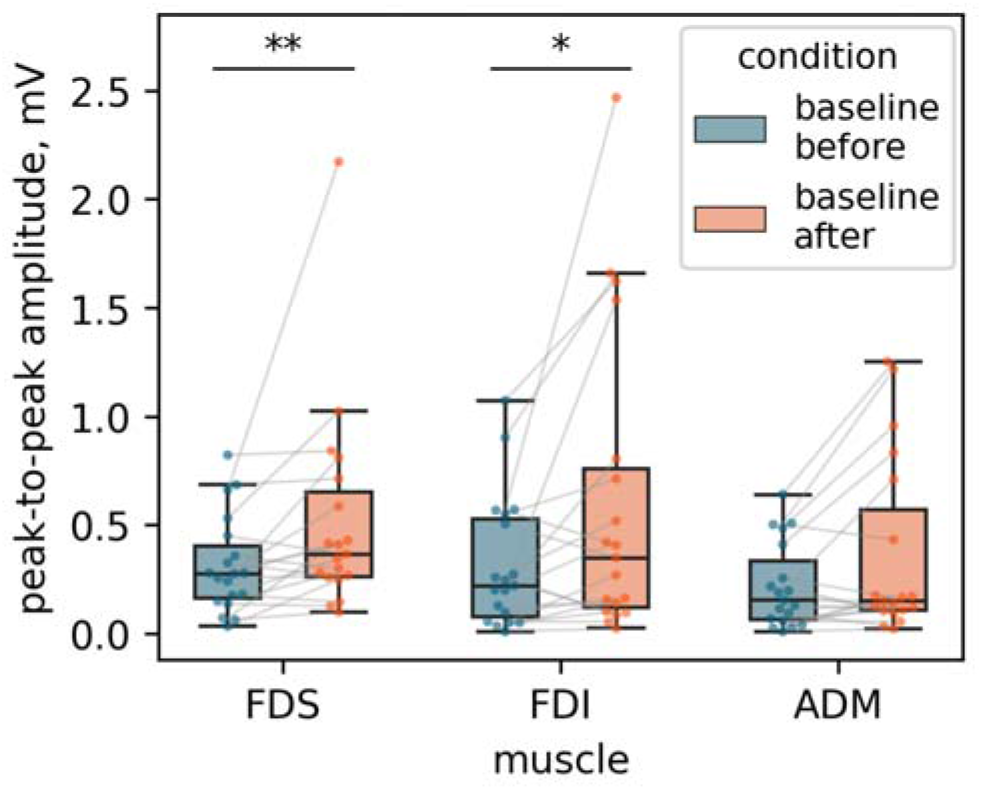
Median peak-to-peak MEP amplitudes for the initial and last baselines for three muscles (*m. flexor digitorum superficialis*, FDS; *m. first dorsal interosseous*, FDI; *m. abductor digiti minimi*, ADM). Values for individual participants are represented by dots. Wilcoxon signed rank test was used for pairwise comparisons where the amplitudes were compared between the initial and final baselines. Asterisks mark statistically significant differences.

## Discussion

The current study adds to the previous reports of the EEG rhythms being modulated by TI (12,16,17). We explored in more detail the differences in effects that TI, kMI and TS exhibited on the TMS-evoked responses in the forearm and hand muscles. In agreement with the numerous previous studies (7,22,23,31), we found that kMI facilitated the MEPs. As such, the kMI condition could serve as a reference to which the effects of TS and TI could be compared. Notably, the effects of TS and TI were weaker than this reference.

Regarding the effects of TS, it affected the MEPs in the FDS and FDI muscles (Fig 2, Fig 3), that is the muscles matching the location of vibratory sensations experienced on the index, middle, and ring fingers, which these muscles move. Although MEPs were weaker during the TS condition as compared to the imagery conditions, no change in MEP was found compared to the baseline. By contrast, some previous studies reported MEP inhibition by TS. Namely, in the experiments of Tokimura et al. (32) peripheral nerve stimulation weakend MEPs when TMS pulse and nerve stimulation were timed to stimulate the motor cortex simultaneously. Such suppression of MEP by a somatosensory afferent input is called short-latency afferent inhibition (SAI). Additionally, in an experimental paradigm that was similar to ours, Oda et al. (33) showed that TS of the right middle fingertip suppressed MEPs in the right FDI. The authors explained this finding in terms of muscle-specific surround inhibition, called sensorimotor gating, where cortical areas representing non-stimulated skin are inhibited to enhance stimulus processing. Furthermore, Kojima et al. (34) showed that TS could modulate corticospinal excitability and that its effects depended on the stimulation patterns. Finally, the findings of Tanaka et al. (35) are similar to our present results: TS applied to the first and second digits did not have an effect on the MEP amplitude in the FDI and ADM. The TMS timing with respect to TS onset being important for MEP inhibition to occur, the absence of this effect in our study could be related to the properties of the stimulation patterns that we used: 100-ms vibratory trains separated by interstimulus intervals of random duration, delivered for 5-8 s. Such a prolonged stimulus is different from the stimuli previously used to achieve SAI.

One of the factors that could have impacted corticospinal excitability during TS is the accommodation of somatosensory receptors caused by the repeated application of vibration. In this study, we made our best effort to minimize any uncontrolled fluctuations in corticospinal excitability and skin sensation. For this purpose, the order of conditions was randomized within a session. The vibro-motor was carefully attached to the skin of the fingertip to stimulate only its surface without applying pressure. Finally, the tactile stimuli were randomly varied in intensity and duration during the TS condition, which mitigated the accommodation and adaptation of somatosensory receptors. The experimental procedures were conducted in a room maintained at a constant temperature. The comparison of MEP amplitudes at the beginning and end of the TS runs did not show any significant differences. We therefore suggest that receptor adaptation did not affect the results for the TS condition.

The comparison of the effects of kMI and TI revealed that kMI significantly increased corticospinal excitability whereas TI had a weaker effect. The observed tendency for different imagery strategies to specifically affect corticospinal excitability agrees with the results of the previous study using F-wave characteristics, where kMI was the type of imagery that enhanced spinal excitability most strongly (26). The modest yet statistically significant increase in the MEPs during TI in our study could have been related to the orientation of attention towards the hand, which agrees with the other studies of the effects of attention to hand position on MEPs amplitude (36).

We found a strong across-subject correlation between the MEP amplitudes during kMI and TI. One possible explanation of this finding is that individual subjects performed kMI and TI in somewhat similar ways, that is partially focused on the muscles even during TI. Such imagery strategy makes given that vibratory stimulation affected not only skin receptors but also muscle spindle afferents (37,38), which could have led to intermixed sensations, even though the participants were instructed to focus on the cutaneous sensations and base their imagery on these sensations, as well. Variability in imagery skills across the participants could also have played a role, with some people possibly relying on the kinesthetic imagery during the TI task. Additionally, the degree to which imagery of any kind (be it TI or kMI) affected corticospinal circuits could have been different across the participants, with some participants resisting these effects and the others being more prone to them.

Finally, we observed a statistically significant difference between the baseline MEPs recorded in the beginning of the experiment and at its end (Fig 4). It is well known that kMI training results in a sustained increase in motor cortex excitability (36), which could have been the case in our study, as well. This post-effect was only statistically significant for FDS and FDI, but not for ADM. We suggest that this is related to the fact that only these muscles were involved in the imagery tasks.

Despite the clear evidence provided by the results of the current study that different types of somatosensory imagery lead to distinct changes in corticospinal excitability, as measured by single-pulse TMS applied over the M1, it is important to acknowledge several limitations of our work.

First and foremost, the effects of the vibratory stimulation utilized in this work are not limited to the activation of cutaneous receptors. Such vibration activates muscle receptors, as well, so it is challenging to produce purely tactile perceptions with such stimulus. To mitigate this disadvantage, alternative stimulation methods, including brush stimulation, von Frey filaments, and pressure-based tactile stimulators, could be employed.

Secondly, we assessed corticospinal excitability by measuring MEPs produced with the TMS coil placed over M1. This stimulation protocol could be enriched by adding different types of stimulation applied to the cortex (e.g., paired-pulse TMS) the spinal cord (e.g., cervical magnetic stimulation) and peripheral nerves (H-reflex, F-wave). These additional methods could contribute to more robust conclusions regarding the effects of TI on corticospinal excitability.

Lastly, for imagery research, additional psychometric measurements could be conducted to assess imagery ability in different domains (e.g., visual, motor, tactile) using specialized scales.

Overall, our study has demonstrated that single-pulse TMS of M1 is specific enough to dissociate the effects of TI and kMI on corticospinal excitability. This finding improves our understanding of the mechanisms of mental imagery, especially in the context of sensorimotor integration and paves way to the imagery-based BCI applications, where kMI and TI exert different effects on the activity of sensorimotor circuits. Such BCI implementations could be used for monitoring and manipulating corticospinal excitability in order to rehabilitate sensorimotor functions in patients suffering from the consequences of stroke and other neural disorders. Distinguishing between different types of somatosensory mental imagery can help to guide the rehabilitation procedures more precisely by accounting for the specifics of brain damage and the affected sensorimotor functions.

## Data Availability

The dataset collected and analyzed during the current study is available at the link https://drive.google.com/drive/folders/1EUW58M5QX9N8uIBNrEJqx2S7l0qjajRe?usp=sharing.

## Competing interests

Authors report no conflict of interest.

## Funding Statement

Funding: MM, AN, LY, NS, and AK were supported by the Russian Foundation, grant #21-75-30024 (https://rscf.ru/en/). This work/article is an output of a research project implemented as part of the Research at the Skolkovo Institute of Science and Technology (Skoltech). The funders had no role in study design, data collection and analysis, decision to publish, or preparation of the manuscript.

## Author contributions

MM, AN, LY and NS collected the data. MM analyzed the data. MM, AN, LY, NS and ML wrote the manuscript. LY, AK and ML supervised the study.

## Notes

https://drive.google.com/drive/folders/1EUW58M5QX9N8uIBNrEJqx2S7l0qjajRe?usp=sharing

### Competing Interest Statement

The authors have declared no competing interest.

### Summary of Updates

Methods section updated to clarify; Discussion improved

## References

1. Gerardin E, Sirigu A, Lehéricy S, Poline JB, Gaymard B, Marsault C, et al. Partially overlapping neural networks for real and imagined hand movements. Cereb Cortex N Y N 1991. 2000 Nov;10(11):1093–104.

2. Guillot A, Collet C, Nguyen VA, Malouin F, Richards C, Doyon J. Brain activity during visual versus kinesthetic imagery: an fMRI study. Hum Brain Mapp. 2009 Jul;30(7):2157–72.

3. Kilintari M, Narayana S, Babajani-Feremi A, Rezaie R, Papanicolaou AC. Brain activation profiles during kinesthetic and visual imagery: An fMRI study. Brain Res. 2016 Sep 1;1646:249–61.

4. Ridderinkhof KR, Brass M. How Kinesthetic Motor Imagery works: a predictive-processing theory of visualization in sports and motor expertise. J Physiol Paris. 2015;109(1–3):53–63.

5. Dickstein R, Deutsch JE. Motor imagery in physical therapist practice. Phys Ther. 2007 Jul;87(7):942–53.

6. García Carrasco D, Aboitiz Cantalapiedra J. Effectiveness of motor imagery or mental practice in functional recovery after stroke: a systematic review. Neurol Barc Spain. 2016;31(1):43–52.

7. Stinear CM, Byblow WD, Steyvers M, Levin O, Swinnen SP. Kinesthetic, but not visual, motor imagery modulates corticomotor excitability. Exp Brain Res. 2006 Jan;168(1– 2):157–64.

8. Toussaint L, Blandin Y. Behavioral evidence for motor imagery ability on position sense improvement following motor imagery practice. Mov Sport Sci - Sci Mot. 2013;(82):63–8.

9. Yang YJ, Jeon EJ, Kim JS, Chung CK. Characterization of kinesthetic motor imagery compared with visual motor imageries. Sci Rep. 2021 Feb 12;11(1):3751.

10. Yoo SS, Freeman DK, McCarthy JJ, Jolesz FA. Neural substrates of tactile imagery: a functional MRI study. Neuroreport. 2003 Mar 24;14(4):581–5.

11. Schmidt TT, Ostwald D, Blankenburg F. Imaging tactile imagery: Changes in brain connectivity support perceptual grounding of mental images in primary sensory cortices. NeuroImage. 2014 Sep 1;98:216–24.

12. Yakovlev L, Syrov N, Miroshnikov A, Lebedev M, Kaplan A. Event-Related Desynchronization induced by Tactile Imagery: an EEG Study. eNeuro [Internet]. 2023 May 1 [cited 2023 Jun 2]; Available from: https://www.eneuro.org/content/early/2023/05/30/ENEURO.0455-22.2023

13. Schmidt TT, Blankenburg F. The Somatotopy of Mental Tactile Imagery. Front Hum Neurosci [Internet]. 2019 [cited 2023 May 19];13. Available from: https://www.frontiersin.org/articles/10.3389/fnhum.2019.00010

14. Bashford L, Rosenthal I, Kellis S, Pejsa K, Kramer D, Lee B, et al. The Neurophysiological Representation of Imagined Somatosensory Percepts in Human Cortex. J Neurosci Off J Soc Neurosci. 2021 Mar 10;41(10):2177–85.

15. Chivukula S, Zhang CY, Aflalo T, Jafari M, Pejsa K, Pouratian N, et al. Neural encoding of actual and imagined touch within human posterior parietal cortex. eLife. 2021 Mar 1;10:e61646.

16. Morozova M, Yakovlev L, Syrov N, Perevoznyuk G, Lebedev M, Kaplan A. Tactile Imagery Affects Cortical Responses to Vibrotactile Stimulation of the Fingertip [Internet]. bioRxiv; 2023 [cited 2023 Jul 18]. p. 2023.06.02.543456. Available from: https://www.biorxiv.org/content/10.1101/2023.06.02.543456v1

17. Yao L, Jiang N, Mrachacz-Kersting N, Zhu X, Farina D, Wang Y. Performance Variation of a Somatosensory BCI Based on Imagined Sensation: A Large Population Study. IEEE Trans Neural Syst Rehabil Eng. 2022;30:2486–93.

18. Yao L, Sheng X, Zhang D, Jiang N, Farina D, Zhu X. A BCI System Based on Somatosensory Attentional Orientation. IEEE Trans Neural Syst Rehabil Eng. 2017 Jan;25(1):81–90.

19. Yao L, Chen ML, Sheng X, Mrachacz-Kersting N, Zhu X, Farina D, et al. A Multi-Class Tactile Brain–Computer Interface Based on Stimulus-Induced Oscillatory Dynamics. IEEE Trans Neural Syst Rehabil Eng. 2018 Jan;26(1):3–10.

20. Kaplan A, Vasilyev A, Liburkina S, Yakovlev L. Poor BCI Performers Still Could Benefit from Motor Imagery Training. In: Schmorrow DD, Fidopiastis CM, editors. Foundations of Augmented Cognition: Neuroergonomics and Operational Neuroscience [Internet]. Cham: Springer International Publishing; 2016 [cited 2023 Jul 18]. p. 46–56. (Lecture Notes in Computer Science; vol. 9743). Available from: http://link.springer.com/10.1007/978-3-319-39955-3_5

21. Vasilyev A, Liburkina S, Yakovlev L, Perepelkina O, Kaplan A. Assessing motor imagery in brain-computer interface training: Psychological and neurophysiological correlates. Neuropsychologia. 2017 Mar;97:56–65.

22. Fadiga L, Buccino G, Craighero L, Fogassi L, Gallese V, Pavesi G. Corticospinal excitability is specifically modulated by motor imagery: A magnetic stimulation study. Neuropsychologia. 1998 Nov;37(2):147–58.

23. Hashimoto R, Rothwell JC. Dynamic changes in corticospinal excitability during motor imagery. Exp Brain Res. 1999 Mar;125(1):75–81.

24. Cengiz B, VurallI D, Zinnuroğlu M, Bayer G, Golmohammadzadeh H, Günendi Z, et al. Analysis of mirror neuron system activation during action observation alone and action observation with motor imagery tasks. Exp Brain Res. 2018 Feb;236(2):497–503.

25. Wright DJ, Wood G, Eaves DL, Bruton AM, Frank C, Franklin ZC. Corticospinal excitability is facilitated by combined action observation and motor imagery of a basketball free throw. Psychol Sport Exerc. 2018 Nov 1;39:114–21.

26. Bunno Y. Imagery strategy affects spinal motor neuron excitability: using kinesthetic and somatosensory imagery. Neuroreport. 2019 May 1;30(7):463–7.

27. Rossi S, Hallett M, Rossini PM, Pascual-Leone A. Safety, ethical considerations, and application guidelines for the use of transcranial magnetic stimulation in clinical practice and research. Clin Neurophysiol. 2009 Dec 1;120(12):2008–39.

28. Pernet C, Garrido MI, Gramfort A, Maurits N, Michel CM, Pang E, et al. Issues and recommendations from the OHBM COBIDAS MEEG committee for reproducible EEG and MEG research. Nat Neurosci. 2020 Dec;23(12):1473–83.

29. Rossini PM, Barker AT, Berardelli A, Caramia MD, Caruso G, Cracco RQ, et al. Non-invasive electrical and magnetic stimulation of the brain, spinal cord and roots: basic principles and procedures for routine clinical application. Report of an IFCN committee. Electroencephalogr Clin Neurophysiol. 1994 Aug;91(2):79–92.

30. Rossini PM, Burke D, Chen R, Cohen LG, Daskalakis Z, Di Iorio R, et al. Non-invasive electrical and magnetic stimulation of the brain, spinal cord, roots and peripheral nerves: Basic principles and procedures for routine clinical and research application. An updated report from an I.F.C.N. Committee. Clin Neurophysiol Off J Int Fed Clin Neurophysiol. 2015 Jun;126(6):1071–107.

31. Kasai T, Kawai S, Kawanishi M, Yahagi S. Evidence for facilitation of motor evoked potentials (MEPs) induced by motor imagery. Brain Res. 1997 Jan 2;744(1):147–50.

32. Tokimura H, Di Lazzaro V, Tokimura Y, Oliviero A, Profice P, Insola A, et al. Short latency inhibition of human hand motor cortex by somatosensory input from the hand. J Physiol. 2000 Mar 1;523(Pt 2):503–13.

33. Oda H, Tsujinaka R, Fukuda S, Sawaguchi Y, Hiraoka K. Tactile Perception of Right Middle Fingertip Suppresses Excitability of Motor Cortex Supplying Right First Dorsal Interosseous Muscle. Neuroscience. 2022 Jul 1;494:82–93.

34. Kojima S, Miyaguchi S, Sasaki R, Tsuiki S, Saito K, Inukai Y, et al. The effects of mechanical tactile stimulation on corticospinal excitability and motor function depend on pin protrusion patterns. Sci Rep. 2019 Nov 13;9(1):16677.

35. Tanaka M, Kubota S, Onmyoji Y, Hirano M, Uehara K, Morishita T, et al. Effect of tactile stimulation on primary motor cortex excitability during action observation combined with motor imagery. Neurosci Lett. 2015 Jul 23;600:1–5.

36. Leung MCM, Spittle M, Kidgell DJ. Corticospinal Excitability Following Short-Term Motor Imagery Training of a Strength Task. J Imag Res Sport Phys Act. 2013 Jun 25;8(1):35–44.

37. Roll JP, Vedel JP. Kinaesthetic role of muscle afferents in man, studied by tendon vibration and microneurography. Exp Brain Res. 1982;47(2):177–90.

38. Cordo P, Gurfinkel VS, Bevan L, Kerr GK. Proprioceptive consequences of tendon vibration during movement. J Neurophysiol. 1995 Oct;74(4):1675–88.

